# Hbxip (Lamtor5) is essential for embryogenesis and regulates embryonic stem cell differentiation through activating mTORC1

**DOI:** 10.1101/2022.01.04.474860

**Authors:** Yan Qin, Peiling Ni, Qingye Zhang, Xiao Wang, Xiaoling Du, Zixi Yin, Lingling Wang, Lihong Ye, Lingyi Chen

## Abstract

Hbxip, also named Lamtor5, has been well characterized as a transcriptional coactivator in various cancers. However, the role of Hbxip in normal development remains unexplored. Here, we demonstrated that homozygous knockout of *Hbxip* leads to embryonic lethality, with retarded growth around E7.5. Using *Hbxip* knockout embryonic stem cells (ESCs), we showed that depletion of Hbxip compromises the self-renewal of ESCs, with reduced expression of pluripotency genes, reduced cell proliferation, and decreased colony forming capacity. In addition, *Hbxip*^-/-^ ESCs are defective in differentiation, particularly ectodermal and mesodermal differentiation. Consistently, *Hbxip*^-/-^ epiblast fails to differentiate properly, indicated by sustained expression of Oct4 in E8.5 *Hbxip*^-/-^ epiblast. Mechanistically, in ESCs, Hbxip interacts with other components of the Ragulator complex, which is required for mTORC1 activation by amino acids. Importantly, ESCs depleted of Ragulator subunits, Lamtor3 or Lamtor4, display differentiation defects similar to those of *Hbxip*^-/-^ ESCs. Moreover, *Hbxip*^-/-^, *p14*^-/-^, and *p18*^-/-^ mice, lacking subunits of the Ragulator complex, also share similar phenotypes, embryonic lethality and retarded growth around E7-8. Thus, we conclude that Hbxip plays a pivotal role in the development and differentiation of the epiblast, as well as the self-renewal and differentiation of ESCs, through activating mTORC1 signaling.

## Introduction

Hepatitis B X-interacting protein (HBXIP, also known as LAMTOR5), was first identified as a binding factor to Hepatitis B virus X protein (Melegari et al., 1998). In addition to its role in inhibiting the replication of Hepatitis B virus, HBXIP has been extensively studied in various cancers. It has been shown that HBXIP promotes proliferation, migration and tumorigenesis of breast, ovarian, liver, non-small-cell lung, bladder urothelial, and esophageal squamous cell cancer (Hu et al., 2011, Liu et al., 2012, Liu et al., 2014, Xu et al., 2014, Zhang et al., 2014, Li and Liu, 2016, Shi et al., 2016, Wang et al., 2017b, Zhou et al., 2019b, Wu et al., 2020). Overexpression of HBXIP is associated with poor prognosis in breast, ovarian, liver, non-small-cell lung cancer and pancreatic ductal adenocarcinomas (Wang et al., 2017b, Li et al., 2017, Wang et al., 2017c, Zhou et al., 2019a, Cheng et al., 2014). HBXIP also contributes to cisplatin and tamoxifen resistance in ovarian and breast cancers, respectively (Zou et al., 2017, Liu et al., 2018). However, the role of HBXIP in normal development remains poorly understood. Only a recent work reported that due to the pivotal role of Hbxip in activating the transcription of insulin by elevating the level of Pdx-1/Neurod1 complex, pancreatic β-cell-specific *Hbxip* knockout (KO) mice display higher fasting blood glucose levels and impaired glucose tolerance (Li et al., 2018).

The extensive studies of HBXIP in cancers have revealed its function as a transcriptional cofactor. HBXIP, cooperating with various transcription factors, such as c-MYB, SP1, cAMP response element-binding protein (CREB), TATA-binding protein (TBP), and E2F1, activates the expression of its downstream target genes, including *YAP, FGF8, LIN28B, SKP2, PDGFB, S100A4*, and *LMO4*, consequently promoting proliferation and migration of cancer cells (Liu et al., 2012, Liu et al., 2014, Xu et al., 2014, Liu et al., 2013, Xu et al., 2013, Yue et al., 2013, Zhang et al., 2013, Wang et al., 2017a). In addition, HBXIP may interact with proteins other than transcription factors to fulfill its biological functions. HBXIP interacts with survivin to suppress caspase activation and hence apoptosis (Marusawa et al., 2003). HBXIP also associates with microtubules and centrosomes in dividing cells, and is required for the proper formation of centrosomes and spindles in HeLa cells (Wen et al., 2008, Fujii et al., 2006). Moreover, HBXIP, together with p18 (LAMTOR1), p14 (LAMTOR2), MP1 (LAMTOR3) and C7ORF59 (LAMTOR4), form a pentameric Ragulator complex, which is required for mTORC1 activation by amino acids (Bar-Peled et al., 2012). Yet, whether HBXIP regulates normal development or carcinogenesis through the mTORC1 pathway remains unexplored.

In this study, we demonstrated that KO of *Hbxip* is embryonic lethal, and retarded development of *Hbxip*^-/-^ embryos become obvious around E7.5. Using *Hbxip* KO embryonic stem cells (ESCs) as an *in vitro* model, we found that *Hbxip* is critical for the differentiation of ESCs, particularly for the ectodermal and mesodermal differentiation. Consistently, the epiblast of E8.5 *Hbxip*^-/-^ embryos remains undifferentiated. The differentiation defects are shared by ESCs lacking subunits of the Ragulator complex, including Hbxip, Lamtor3, and Lamtor4. Thus, we concluded that Hbxip regulates embryo development and ESC differentiation through activating the mTORC1 signaling.

## Results

### Embryonic lethality in *Hbxip* null embryos

To study the function of Hbxip in normal development, we knocked out *Hbxip* in previous generated *Hbxip* floxed mice (Figure 1A) (Li et al., 2018). Heterozygous *Hbxip* KO (*Hbxip*^+/-^) mice were obtained through mating between *Hbxip* floxed mice and *EIIa-cre* mice. *Hbxip*^+/-^ mice are fertile. Yet, no *Hbxip*^-/-^ mice were born from *Hbxip*^*+/-*^ intercrosses (Figure 1C), indicating embryonic lethality of *Hbxip* null mice. To determine the timing of lethality, E3.5, E6.5, E7.5 and E9.5 embryos from *Hbxip*^*+/-*^ intercrosses were genotyped, and *Hbxip*^-/-^ embryos were detected at all the analyzed time points (Figure 1C). However, at 7.5 days postcoitum, *Hbxip*^-/-^ embryos are smaller than WT and *Hbxip*^*+/-*^ embryos (Figure 1D and E). At 8.5 and 9.5 days postcoitum, *Hbxip*^-/-^ embryos appear to be developmentally arrested, compared to their WT and *Hbxip*^*+/-*^ littermates (Figure 1D). These data suggest a critical role of Hbxip in embryonic development.

**Figure 1.**
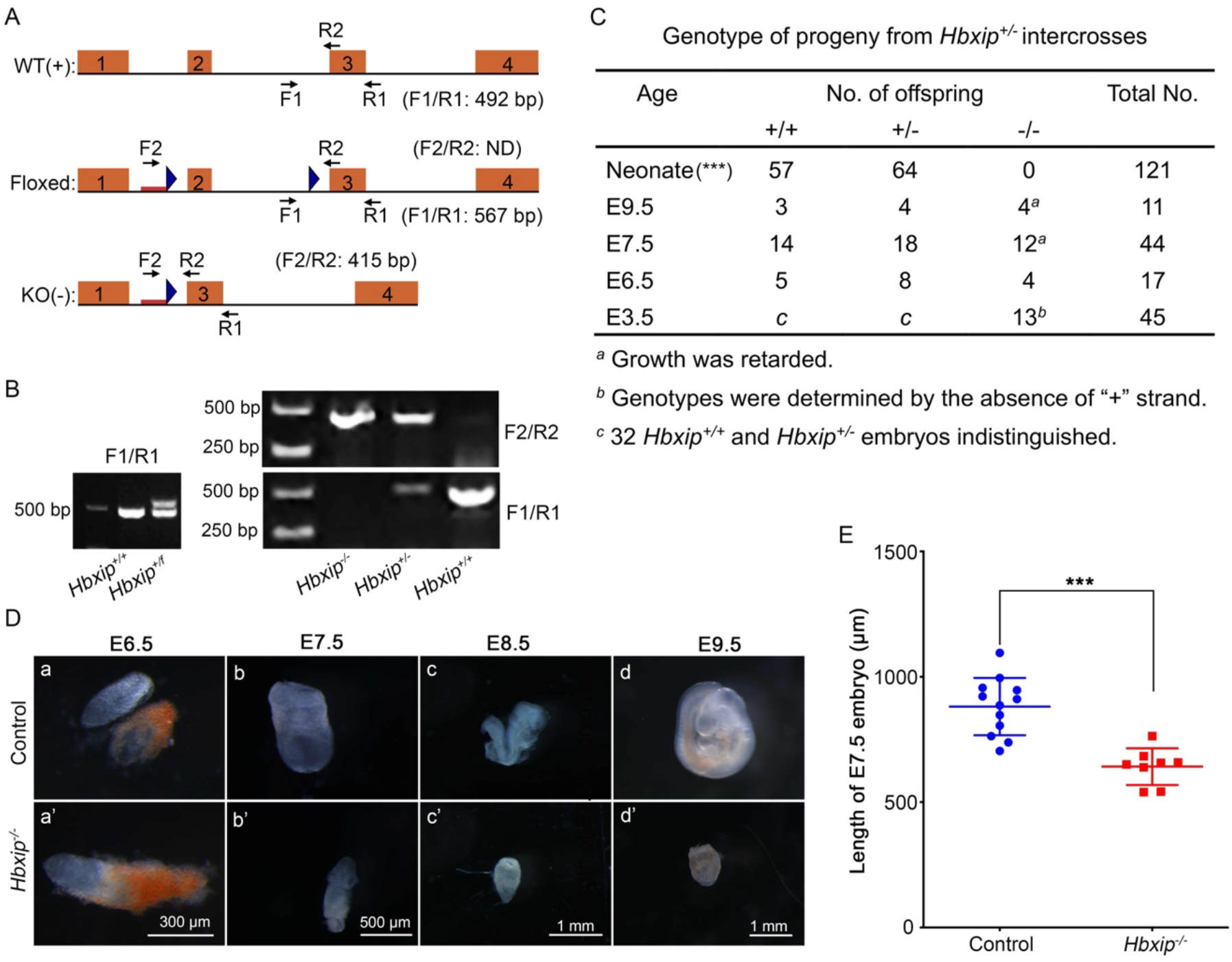
Embryonic lethality in *Hbxip*^*-/-*^ mice. (A) Schematic illustration of WT, floxed, and KO alleles of *Hbxip*. Orange rectangles represent *Hbxip* exons, and blue triangles marks loxP sequences. Red bars in the floxed and KO alleles are the exogenous DNA fragment from the targeting vector. Arrows show the primers for genotyping PCR. (B) Representative genotyping PCR results. (C) Genotyping of E3.5, E6.5, E7.5, E9.5 embryos and neonate mice derived from the intercrossing of *Hbxip*^*+/-*^ mice. *χ*^2^ test was performed on the number of mouse mutants obtained per stage in comparison to the expected Mendelian ratios. ***, p<0.001. (D) The morphology of dissected control (including WT and *Hbxip*^+/-^) and *Hbxip*^-/-^ embryos, without any staining, at various stages. (a) E6.5 Control, n=13; (a’) E6.5 *Hbxip*^-/-^, n=4; (b) E7.5 Control, n=12; (b’) E7.5 *Hbxip*^-/-^, n=8; (c) E8.5 Control, n=8; (c’) E8.5 *Hbxip*^-/-^, n=6; (d) E9.5 Control, n=7; (d’) E9.5 *Hbxip*^-/-^, n=4. (E) Quantification of the length of E7.5 *Hbxip*^*-/-*^ and control (including WT and *Hbxip*^+/-^) embryos. Control, n=12; *Hbxip*^-/-^, n=8. Statistical analysis was performed with Student’s *t* test. ***, p<0.001.

### *Hbxip* KO compromises the self-renewal of embryonic stem cells

Next, we utilized *in vitro* cultured mouse embryonic stem cells (ESCs) to understand the mechanism of Hbxip in embryonic development. *Hbxip* KO ESC lines were constructed using CRISPR/Cas9, with a sgRNA targeting to the third exon of *Hbxip* (Figure 2A). The disruption of *Hbxip* in two independent ESC clones (hereafter referred to as *H*^-/-^*-1* and *H*^-/-^*-2*) was validated by loss of the *Ban* I site, Sanger sequencing, and Western Blot (Figure 2A, B and S1). Notably, deletion of *Hbxip* compromises the self-renewal of ESCs, demonstrated by reduced proliferation rate and colony-forming capacity (Figure 2C and D). The mRNA levels of pluripotency genes, *Nanog, Oct4*, and *Sox2*, as well as the protein levels of Nanog and Oct4, are declined in *Hbxip* KO ESCs (Figure 2B and E). In addition, in undifferentiated ESCs, *Hbxip* KO suppresses the expression of ectodermal, and mesodermal markers, such as *Nestin, Celsr2, T*, and *Dlx3*, whereas endodermal marker *Gata6* is activated by *Hbxip* KO (Figure 2F). All these data suggest an essential role of *Hbxip* in ESC self-renewal.

**Figure 2.**
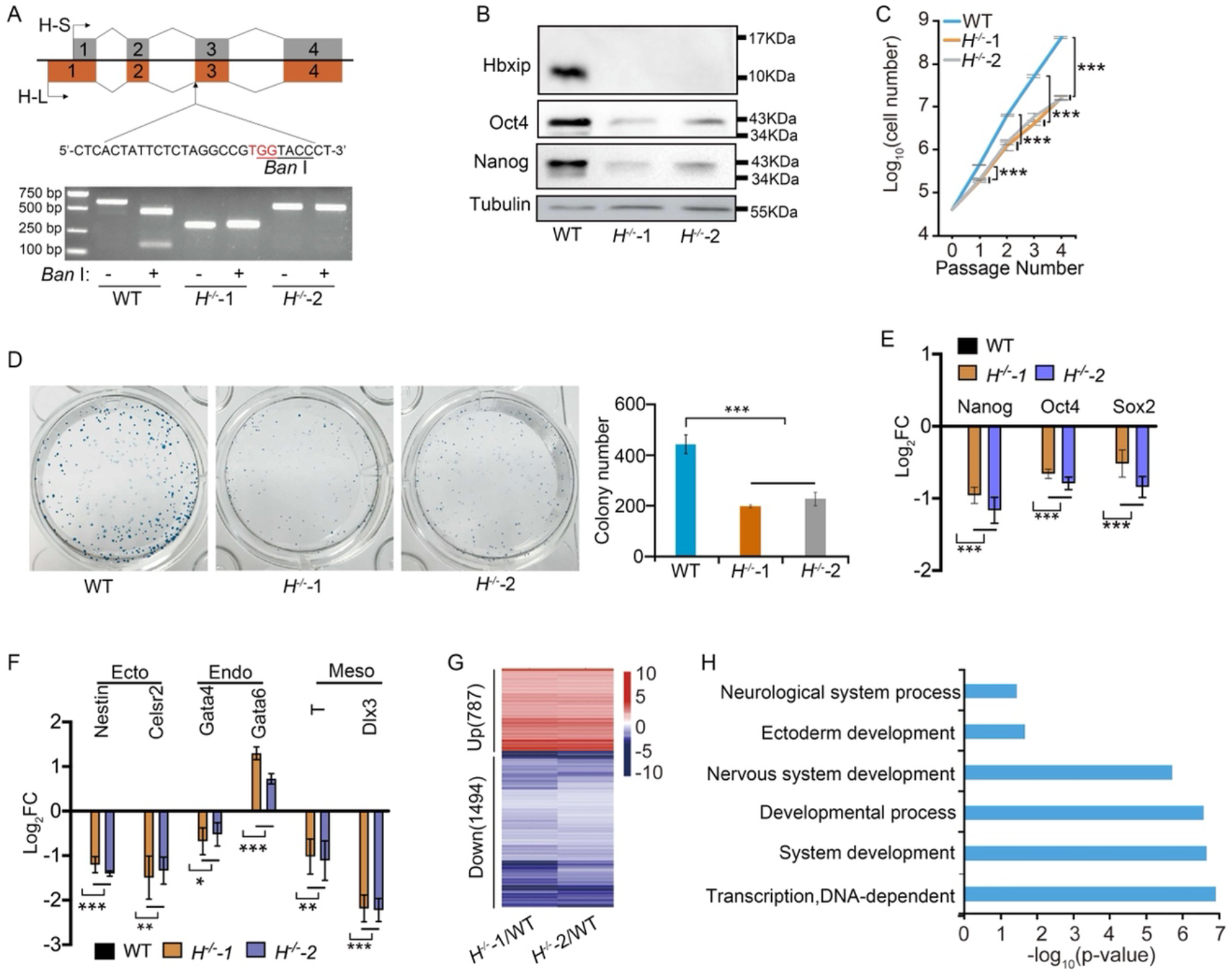
Compromised self-renewal of *Hbxip*^*-/-*^ ESCs. (A) Schematic illustration of the strategy for knocking out *Hbxip* in ESCs. The exons of two *Hbxip* isoforms, H-S and H-L, are represented by grey and orange rectangles, respectively. The targeting sequence of sgRNA is shown. The protospacer-adjacent motif (PAM) is highlighted in red, and the *Ban* I site is underlined. The bottom panel shows the validation of *Hbxip* KO in two independent ESC clones by PCR and *Ban* I digestion. (B) Western blots show that the levels of pluripotency factors Nanog and Oct4 are reduced in *Hbxip*^-/-^ ESCs. (C) Growth curves of WT and *Hbxip*^-/-^ ESCs. (D) Colony forming assay of WT and *Hbxip*^-/-^ ESCs. The left panel shows representative AP staining images of colony forming assays. The right panel is the quantification results of three repeated colony forming assays. (E) The expression of pluripotency genes *Nanog, Oct4*, and *Sox2* in WT and *Hbxip*^-/-^ ESCs, measured by quantitative RT-PCR. Fold change (FC) was calculated by comparing to WT ESCs. (F) The expression of differentiation genes in WT and *Hbxip*^-/-^ ESCs. FC was calculated by comparing to WT ESCs. (G) The heatmap showing the differentially expressed genes in *H*^*-/*-^-1 and *H*^*-/*^-2 ESCs, compared to WT ESCs, detected by RNA-seq. (H) GO annotation of the downregulated genes in *Hbxip*^-/-^ ESCs. For growth curve, colony formation, quantitative RT-PCR and Western blot, n=3. Statistical analysis was performed with two-way ANOVA test. *, p < 0.05; **, p < 0.01; ***, p < 0.001.

Transcriptomic analysis identified 787 upregulated genes and 1494 downregulated genes shared by *H*^-/-^*-1* and *H*^-/-^*-2* ESCs, compared to WT ESCs (Figure 2G, and Table S2). Gene ontology (GO) annotation revealed that several developmental terms, such as system development, nervous system development and ectoderm development, are enriched in the downregulated genes (Figure 2H and S2A). These data imply that Hbxip might be involved in ESC and epiblast differentiation.

### *Hbxip* KO causes differentiation defects in ESCs

We then tested the role of Hbxip in the differentiation of ESCs. Two methods were used to differentiate ESCs, either by LIF withdrawal or by embryoid body (EB) formation in hanging drops. Under these two differentiation conditions, *Hbxip*^-/-^ ESCs fail to activate differentiation genes, including ectodermal markers, *Nestin* and *Celsr2*, and mesodermal markers, *T* and *Dlx3* (Figure 3A and B). Through RNA-seq analysis, 1028 downregulated genes and 482 upregulated genes are identified in differentiated *Hbxip*^-/-^ cells induced by LIF withdrawal, compared to differentiated WT cells (Figure 3C), while *Hbxip* KO downregulates 719 genes and upregulates 655 genes in day 4 EBs (Figure 3D and Table S3). Differentiated *Hbxip*^-/-^ cells by LIF withdrawal and day 4 *Hbxip*^-/-^ EBs share 481 downregulated genes and 325 upregulated genes (Figure 3E and Table S3). GO analysis revealed that genes involved in system development, mesoderm development, ectoderm development, embryo development and cell differentiation are enriched in the downregulated genes (Figure 3F and S2B).

**Figure 3.**
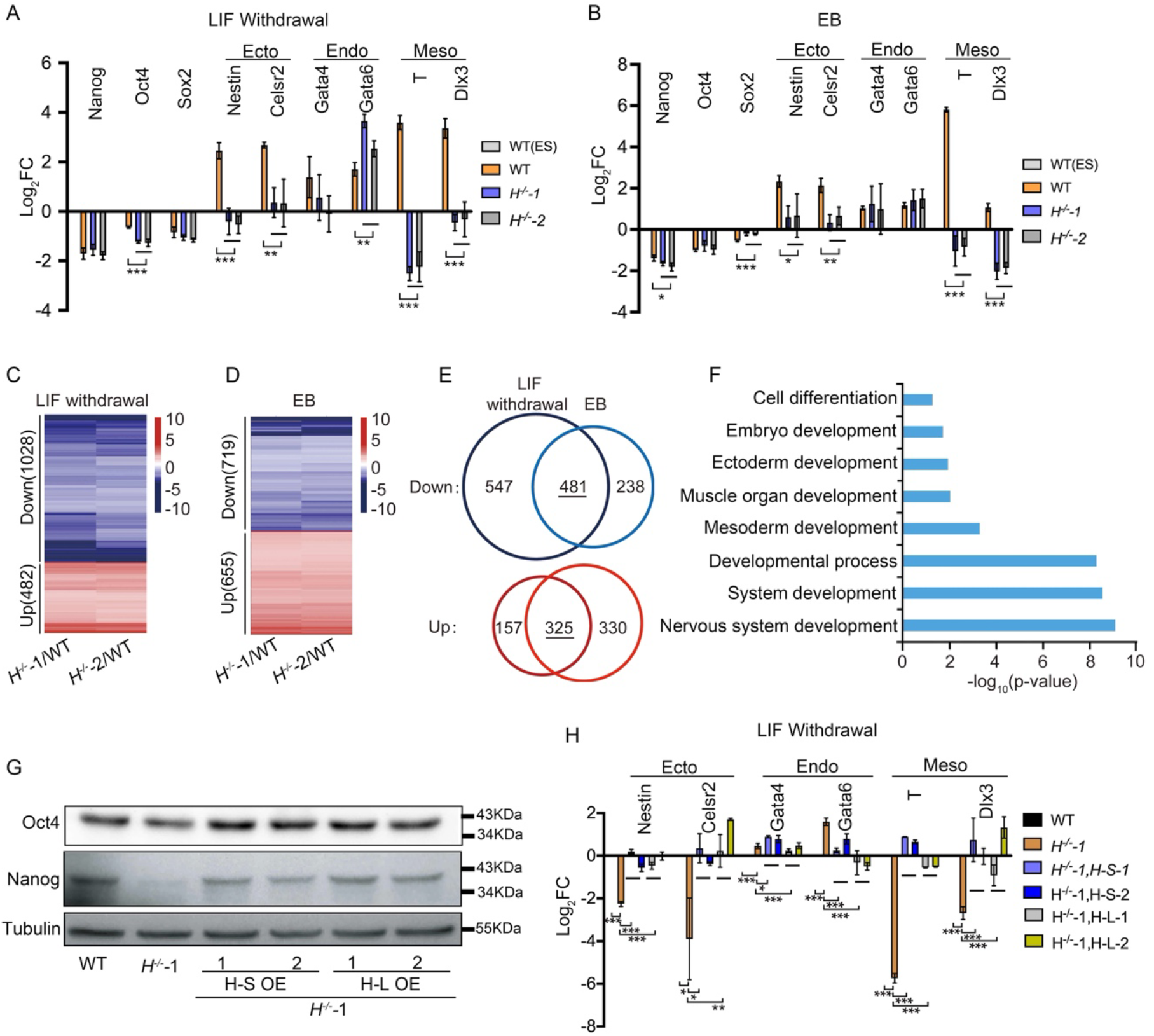
Differentiation defects of *Hbxip*^*-/-*^ ESCs. (A) and (B) The expression of differentiation genes in differentiated WT and *Hbxip*^-/-^ ESCs by LIF-withdrawal for 4 days (A) and in day 4 WT and *Hbxip*^-/-^ EBs (B). FC was calculated by comparing to WT ESCs. (C) and (D) The heatmaps show the differentially expressed genes in differentiated *Hbxip*^-/-^ ESCs by LIF-withdrawal for 4 days (C) and day 4 *Hbxip*^-/-^ EBs (D), compared to their WT counterparts, detected by RNA-seq. (E) Venn diagrams show the commonly regulated genes (481 downregulated and 325 upregulated genes) by *Hbxip* KO in two differentiation protocols, LIF-withdrawal and EB differentiation. (F) GO annotation of the commonly downregulated genes by *Hbxip* KO. (G) Overexpression of *Hbxip*, both H-L and H-S, rescues the expression levels of Nanog and Oct4 proteins in *Hbxip* KO ESCs. (H) Overexpression of *Hbxip*, both H-L and H-S, rescues the expression levels of differentiation genes in differentiated *Hbxip* KO ESCs by LIF-withdrawal for 4 days. FC was calculated by comparing to differentiated WT ESCs. For quantitative RT-PCR and Western blot, n=3. Statistical analysis was performed with two-way ANOVA test. *, p < 0.05; **, p < 0.01; ***, p < 0.001.

To further confirm the roles of *Hbxip* in pluripotency maintenance and ESC differentiation, rescue experiments were performed in *H*^*-/-*^*-1* ESCs by expressing either short isoform (H-S) or long isoform (H-L) of *Hbxip*. Both H-S and H-L rescue the expression levels of Nanog and Oct4 proteins in undifferentiated ESCs (Figure 3G and S2C), as well as the expression of differentiation genes, *Nestin, Clesr, T*, and *Dlx3*, in differentiated cells (Figure 3H). These data indicate that Hbxip is required for the proper differentiation of ESCs, particularly toward the ectodermal and mesodermal lineages.

### *Hbxip*^*-/-*^ embryos are defective in epiblast formation and differentiation

Given the differentiation defects of *Hbxip*^*-/-*^ ESCs, we tested whether the epiblast properly differentiates into three germ layers, particularly the ectoderm and mesoderm. Immunohistochemistry staining with Hbxip antibody allowed us to distinguish *Hbxip*^*-/-*^ embryos from WT and *Hbxip*^*+/-*^ embryos (Figure 4). Hematoxylin-eosin (H&E) staining showed that the development of amnion cavity is retarded in E7.5 *Hbxip*^*-/-*^ embryos (Figure 4A).

**Figure 4.**
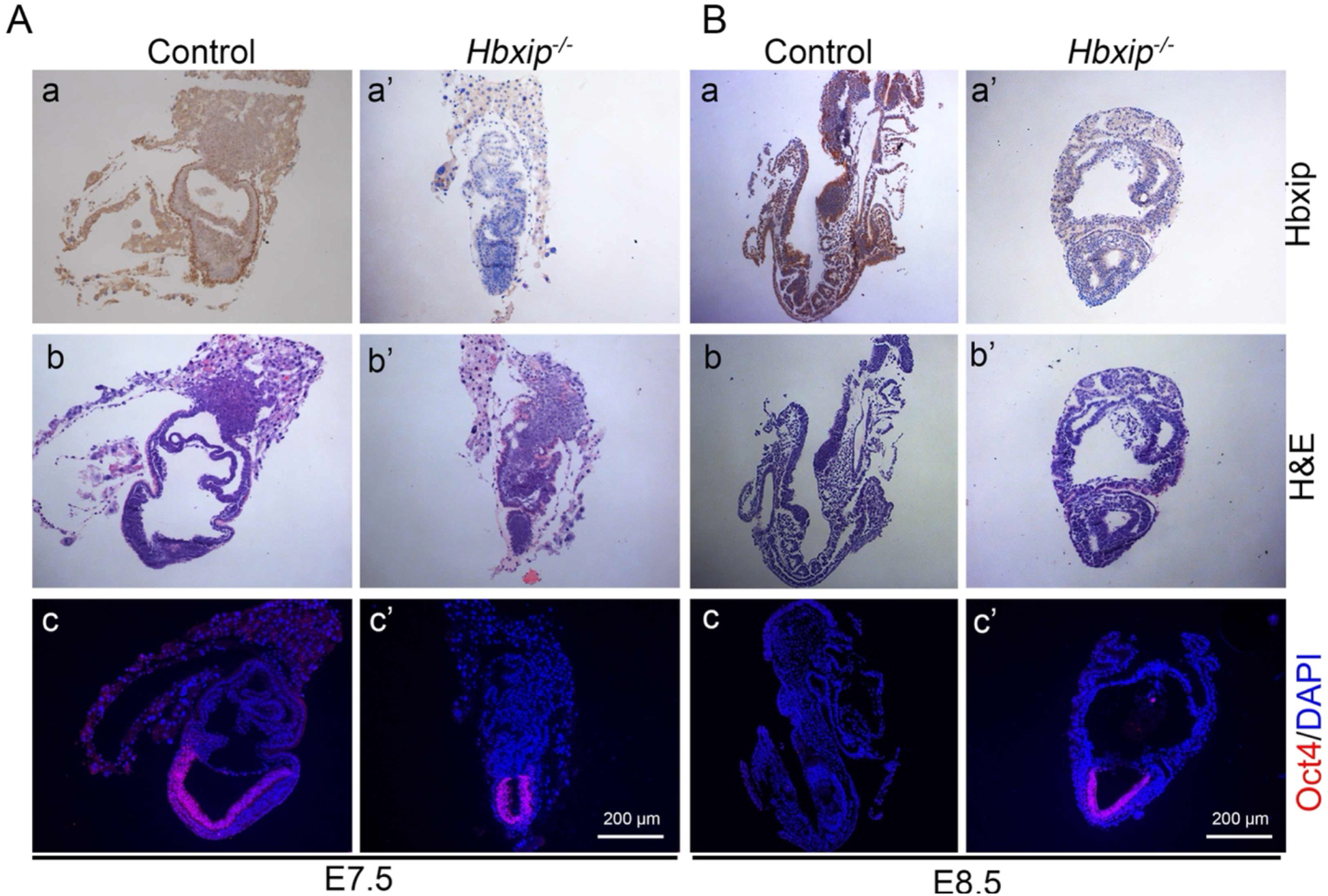
Defective epiblast proliferation and differentiation in *Hbxip*^*-/-*^ embryos. Control and *Hbxip*^*-/-*^ E7.5 (A) and E8.5 (B) embryo sections were subjected to immunohistochemistry staining for Hbxip (a and a’), H&E staining (b and b’), and immunofluorescence staining for Oct4 (c and c’). E7.5 (Control, n=14; *Hbxip*^-/-^, n=8); E8.5 (Control, n=8; *Hbxip*^-/-^, n=5).

Immunofluorescent staining of Oct4 demonstrated that the epiblast in E7.5 *Hbxip*^*-/-*^ embryos is much smaller than that in E7.5 WT and *Hbxip*^*+/-*^ embryos, indicating defective epiblast formation (Figure 4A). At 8.5 days postcoitum, the morphology of WT and *Hbxip*^*+/-*^ embryos changes drastically as the initiation of somitogenesis. In contrast, the development of *Hbxip*^*-/-*^ embryos does not progress further. Importantly, with the formation of three germ layers, Oct4 expression is diminished in E8.5 WT and *Hbxip*^*+/-*^ embryos, whereas *Hbxip*^*-/-*^ embryos retain Oct4 expression in the epiblast (Figure 4B). Taken together, these results suggest that *Hbxip*^*-/-*^ embryo lethality is due to the defects in epiblast proliferation and differentiation.

### Hbxip is required for the activation of mTORC1

Next, we investigated the molecular mechanism of Hbxip in embryonic development and ESC differentiation. It has been shown that HBXIP functions as a transcriptional cofactor in many cancers (Liu et al., 2012, Liu et al., 2014, Xu et al., 2014, Liu et al., 2013, Xu et al., 2013, Yue et al., 2013, Zhang et al., 2013, Wang et al., 2017a). We first determined the subcellular distribution of Hbxip in ECSs by Western blot with cytoplasmic and nuclear fractions of ESCs. The result showed that Hbxip is almost exclusively distributed in the cytoplasm (Figure 5A), thus excluding its function as a transcriptional cofactor.

**Figure 5.**
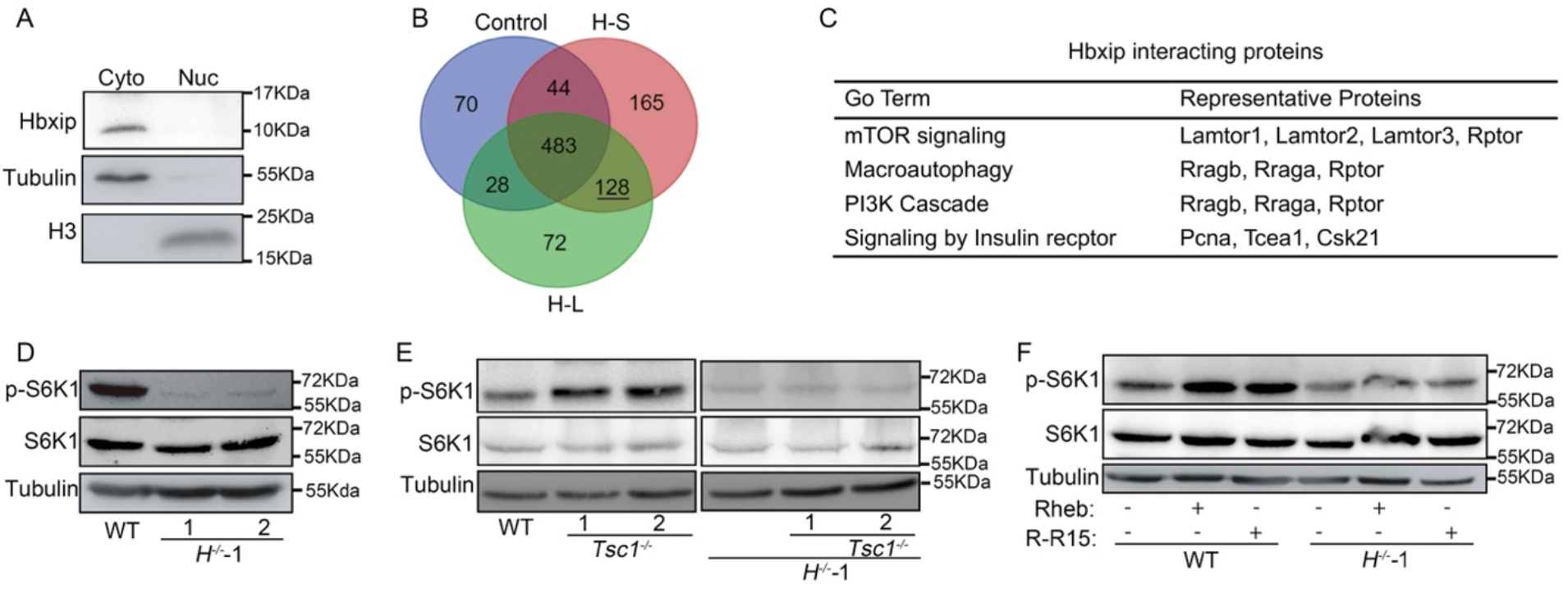
Hbxip is required for mTORC1 activation. (A) Western blot analysis of Hbxip in the cytoplasmic and nuclear fractions of ESCs. (B) Venn diagram of proteins identified by mass spectrometric analysis. Co-IP experiments were performed with cell extracts from ESCs expressing FLAG-Hbxip (H-L and H-S) and control ESCs with empty vector, using M2 magnetic beads. The resulting IP samples were analyzed by mass spectrometry. 128 proteins identified in both the H-L and H-S samples, but not in the control, are called as Hbxip interacting proteins. (C) GO annotation of the 128 Hbxip-interacting proteins. (D) Western blots of S6K1 and p-S6K1 in WT and *Hbxip*^-/-^ ESCs demonstrate the reduced mTORC1 activity in *Hbxip*^-/-^ ESCs. (E) Western blots of S6K1 and p-S6K1 in WT, *Tsc1*^-/-^, *H*^-/-^-1, and *Tsc1*^-/-^; *H*^-/-^-1 ESCs show that *Tsc1* KO fails to activate mTORC1 in *Hbxip*^-/-^ ESCs. (F) Overexpression of Rheb or Rptor fused with the lysosome-targeting signal of Rheb (R-R15) is unable to activate mTORC1 in *Hbxip*^-/-^ ESCs. WT and *H*^-/-^-1 ESCs were transfected with empty vector, or plasmids expressing Rheb or R-R15. Two days after transfection, ESCs were harvested for Western blot analysis. For Western blot, n=3.

To understand the molecular mechanism of Hbxip, coimmunoprecipitation (co-IP) coupled with mass spectrometric analysis was performed to identify Hbxip interacting proteins. 128 proteins were identified in the H-S and H-L co-IP samples, but not in the control co-IP sample (Figure 5B and Table S4). GO analysis revealed that these Hbxip interacting proteins are enriched in mTOR signaling, macroautophagy, PI3K cascade, and signaling by insulin receptor (Figure 5C). It has been shown that HBXIP is a member of the Ragulator complex which recruits mTORC1 to the lysosome, therefore activating the mTORC1 signaling pathway (Bar-Peled et al., 2012). Consistently, three Ragulator components, Lamtor1 (p18), Lamtor2 (p14), and Lamtor3 (MP1), as well as a mTORC1 subunit Rptor, were identified as Hbxip interacting proteins (Figure 5C and Table S4). Thus, we speculated that mTORC1 is the key downstream factor of Hbxip in ESCs and embryos. To test this hypothesis, we first showed that the activity of mTORC1 is reduced in *Hbxip*^*-/-*^ ESCs and embryos, as indicated by the level of phosphorylated S6K1 and 4EBP1 (Figure 5D, S2D, and S3). Next, we tried to reactivate mTORC1 to rescue the differentiation defects of *Hbxip*^*-/-*^ ESCs. However, even though knockout of *Tsc1*, overexpression of Rheb or Rptor fused with the lysosome-targeting signal of Rheb (R-R15) (Sancak et al., 2010, Duvel et al., 2010, Long et al., 2005), successfully activate mTORC1 in WT ESCs, these strategies failed to activate the mTORC1 signaling in *Hbxip*^*-/-*^ ESCs (Figure 5E, 5F and S4), demonstrating the essential role of Hbxip in mTORC1 activation.

### mTORC1 inactivation accounts for the defects in ESC self-renewal and differentiation

Failure in activating mTORC1 in *Hbxip*^*-/-*^ ESCs indicates that Hbxip is essential for mTORC1 activation. Yet, it rendered the rescue experiment impossible. To prove that inactivation of mTORC1 is responsible for the self-renewal and differentiation defects in *Hbxip*^*-/-*^ ESCs, an alternative strategy was applied. Instead of the rescue experiment, we knocked out genes encoding other components of the Ragulator complex, such as *Lamtor3* and *Lamtor4*, in ESCs (Figure S5). Similar to *Hbxip*^-/-^ ESCs, *Lamtor3*^-/-^ and *Lamtor4*^-/-^ ESCs display self-renewal defects, including reduced protein levels of Oct4 and Nanog, down-regulated *Oct4, Nanog*, and *Sox2* mRNA expression, slower proliferation rate, and decreased colony forming capacity (Figure 6A-D). Moreover, upon differentiation, *Lamtor3*^-/-^ and *Lamtor4*^-/-^ ESCs also fail to activate ectodermal and mesodermal markers (Figure 6E and F), just as what we observed in *Hbxip*^-/-^ ESCs (Figure 3A and B). All these data indicate that inactivation of mTORC1 leads to the self-renewal and differentiation defects in ESCs.

**Figure 6.**
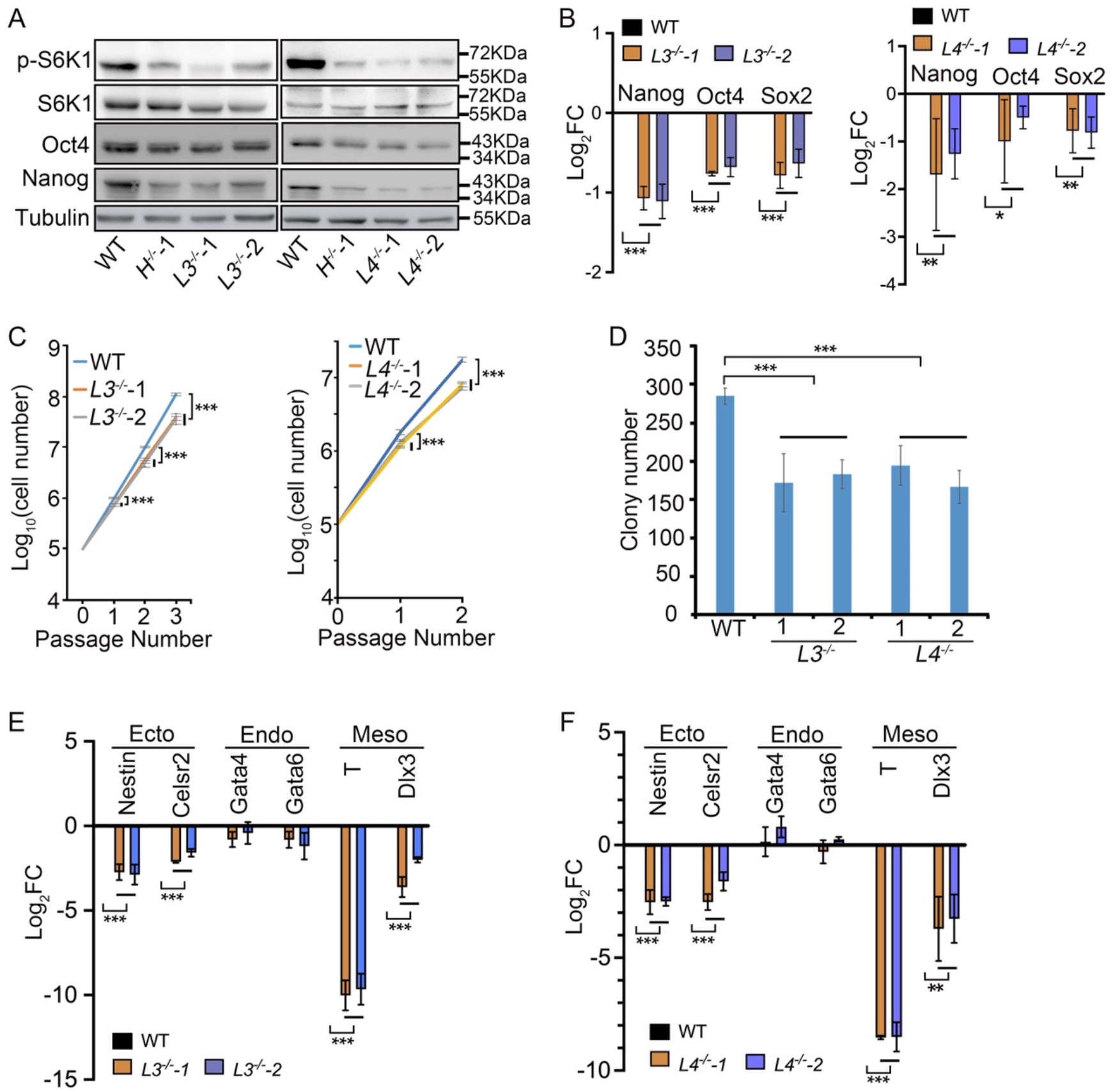
Differentiation defects in ESCs lacking other Ragulator subunits. (A) Western blots of S6K1, p-S6K1, Nanog, and Oct4 in WT, *Hbxip*^-/-^, *Lamtor3*^-/-^ (*L3*^-/-^) and *Lamtor4*^-/-^ (*L4*^-/-^) ESCs. (B) The expression levels of pluripotency genes in WT, *L3*^-/-^ and *L4*^-/-^ ESCs measured by quantitative RT-PCR. FC was calculated by comparing to WT ESCs. (C) Growth curves of WT, *L3*^-/-^ and *L4*^-/-^ ESCs. (D) Colony forming assays of WT, *L3*^-/-^ and *L4*^-/-^ ESCs. (E) and (F) The expression levels of differentiation genes in differentiated WT, *L3*^-/-^ (E) and *L4*^-/-^ (F) ESCs. ESCs were differentiated for 96 hours in the absence of LIF. The resulting differentiated cells were harvested for quantitative RT-PCR. FC was calculated by comparing to differentiated WT ESCs. For growth curve, colony formation, quantitative RT-PCR and Western Blots, n=3. Statistical analysis was performed with two-way ANOVA test. *, p < 0.05; **, p < 0.01; ***, p < 0.001.

## Discussion

In this study, we demonstrated that Hbxip is essential for embryonic development and ESC differentiation. *Hbxip*^-/-^ ESCs, as well as *Lamtor3*^-/-^, and *Lamtor4*^-/-^ ESCs, with disrupted Ragulator complex, display differentiation defects toward ectodermal and mesodermal lineages (Figure 3A 3B, 6E, and 6F). Moreover, *Hbxip*^-/-^, *p14*^-/-^, and *p18*^-/-^ mice, lacking subunits of the Ragulator complex, are embryonic lethal, and retarded growth of these embryos are detected at the same developmental stage, around E7-8 (Figure 1C and D) (Nada et al., 2009, Teis et al., 2006). These data indicate that the phenotypes of *Hbxip*^-/-^ embryos and ESCs are mainly caused by loss function of the Ragulator complex, rather than lacking the transcriptional coactivator role of Hbxip.

The Ragulator complex is required for the activation of mTORC1 by amino acids (Sancak et al., 2010, Bar-Peled et al., 2012). Consistently, the mTORC1 activity is reduced in *Hbxip*^-/-^ ESCs (Figure 5D and S2D). KO of *Tsc1* and overexpression of Rheb or R-R15 fail to activate mTORC1 in the absence of Hbxip (Figure 5E and F). In addition, the reduced mTORC1 activity by disruption of the Ragulator complex allows us to investigate the function of mTORC1 in post-implantation embryo development. Deletion of mTORC1 components, mTor and Rptor, results in peri-implantation embryonic death around E5.5-6.5 (Gangloff et al., 2004, Murakami et al., 2004, Guertin et al., 2006), thus preventing studying the role of mTORC1 in later stage of embryogenesis. Nevertheless, *Hbxip*^-/-^, *p14*^-/-^, and *p18*^-/-^ embryos, in which mTORC1 activity is presumably attenuated, develop further after implantation, and retarded growth appears around E7-8. The reduced size of E7.5 *Hbxip*^-/-^ embryos might be attributed to slower cell proliferation rate caused by decreased mTORC1 activity. Moreover, the differentiation defects of *Hbxip*^-/-^ ESCs (Figure 3A and B) imply that *Hbxip*^-/-^ epiblast might be defective in differentiation. Indeed, from E7.5 to E8.5, *Hbxip*^-/-^ epiblast appears undifferentiated, as indicated by the sustained expression of Oct4 (Figure 4), suggesting that mTORC1 activity is pivotal for epiblast differentiation. Consistent with this note, inhibition of mTOR by INK128 induces a paused pluripotent state in the blastocyst, mimic diapause (Bulut-Karslioglu et al., 2016), further supporting the role of mTORC1 in the exit from pluripotency.

mTOR signaling is involved in growth and disease through regulating gene transcription, mRNA degradation, protein synthesis, autophagy, lipid and carbohydrate metabolism (Giguere, 2018, Villa et al., 2021, Cho et al., 2021, Mossmann et al., 2018, Saxton and Sabatini, 2017). Further studies are required to elucidate how mTORC1 regulated by the Ragulator complex promotes the differentiation of the epiblast and ESCs. *Hbxip*^-/-^, *Lamtor3*^-/-^, and *Lamtor4*^-/-^ ESCs provide an excellent *in vitro* experimental system for mechanistic investigation.

## Materials and Methods

### Mice

Nankai Animal Care and Use Committee approved the use of mice for this research (Approval number: 2021-SYDWLL-000469). *EIIa-cre* mice (Lakso et al., 1996), which express Cre in early mouse embryos and is useful for whole-body and germ line deletion of floxed allele, were purchased from Shanghai Model Organisms. *Hbxip*^-/-^ mice were obtained by crossing *Hbxip*^+/-^ mice. Genotyping was performed as described previously (Qin et al., 2019). DNA was isolated from dissected embryos or the tail tips of 2-week mice, and then analyzed by PCR. The sequences of primers (illustrated in Figure 1A) are listed below. F1: 5’-TTTTTGTCACTCTCGCCTTTG-3’, R1: 5’-GCTGGTATGTACTCACCCATT-3’, F2: 5’-GCTCTATGGCTTCTGAGGCGGAA, R2: 5’-CTCATCCGACAGGGTACCACGG-3’.

### Embryo collection

Embryo manipulation experiments were carried out as described previously (Zhou et al., 2018). Female *Hbxip*^*+/-*^ mice (4-6 weeks) were induced to superovulate by intraperitoneal injection of 5 IU pregnant mare serum gonadotropin (PMSG, Calbiochem) and 48 hours later 5 IU human chorionic gonadotropin (hCG, Sigma). Then females were mated with male *Hbxip*^*+/-*^ mice overnight. The next morning, females were checked for the presence of a vaginal plug, designed as E0.5. Zygote and E3.5 embryos were flushed from oviducts at day 0.5 or day 3.5 post-hCG. E6.5, E7.5, E8.5 and E9.5 embryos were collected at corresponding time points after successful mating.

### Histological analysis

Embryos were dissected from mice immediately after euthanasia, fixed in 4% paraformaldehyde (PFA) for up to 24 h, stored in 70% ethanol and embedded in paraffin. Then 5 μm-thick sections were prepared using rotary microtome (Leica) and mounted on glass slides. After deparaffinization, sections were stained with hematoxylin and eosin (H&E) for histological analysis.

### Immunohistochemistry (IHC) analysis

After deparaffinization, sections were boiled in 1× citrate buffer (ORIGENE, ZLI-9064) and then microwaved at low power for 15 minutes. After cooling to room temperature (RT), the sections were blocked with 5% bovine serum albumin (BSA) at RT for 45 minutes, and then incubated with primary antibody (*α*-Hbxip, CST, 14633S) at RT for 1 h. After three washes with 1×PBS buffer and 3% H2O2 treatment for 10 minutes, the slides were incubated with goat anti-rabbit IgG conjugated to horseradish peroxidase (HRP) (ORIGENE, ZB-2301) at RT for 1 h. The HRP activity was detected by the Vectastain ABC (avidin-biotin-peroxidase) kit (Vector Laboratories) as recommended by the manufacturer. The slides were examined using a Leica Microscope, and images were captured with a Leica DFC420C camera.

### Immunofluorescence (IF) analysis

The IF of paraffin sections was carried out as described previously (Qin et al., 2019). After deparaffinization, the sections were blocked with 5% goat serum at RT for 45 minutes, then incubating with primary antibody (*α*-Oct4, Abcam, ab181557; *α*-p-S6K1, Sigma, SAB4301518) at RT for 1 h. After washing in 1×PBS buffer for three times, the slides were incubated with Alexa Fluor 594 conjugated goat anti-rabbit IgG (Invitrogen, A11037) at RT for 1 h. The nucleus was stained with DAPI (Thermo Fisher). All images were captured with Leica DM3000 microscope with DFC420C camera.

### Cell culture and Transfection

Mouse ESCs (E14 and its derivatives) were cultured in high glucose DMEM (GIBCO) supplemented with 15% fetal bovine serum (FBS, Hyclone), 1% penicillin/streptomycin, 1% L-glutamine, 1% β-mercaptoethanol (Sigma), 1% non-essential amino acids (NEAA, Invitrogen) and 1000 U/mL LIF (ESGRO, Chemicon), plated on gelatin-coated tissue culture dish, and grown in a humidified 5% CO2 at 37°C. Transfection was carried out with Lipofectamine^®^ 3000 (Invitrogen) according to the manufacturer’s instruction. For construction of stably transfected clones, puromycin (1.25 μM) selection was applied for 5-7 days, starting from 24 hours after transfection.

### Vector construction

pX330-U6-Chimeric_BB-CBh-hSpCas9 (pX330) vector was used to construct gene knockout plasmids targeting *Hbxip, Tsc1, Lamtor3* and *Lamtor4*. sgRNA was designed in Zhang Feng’s online website (http://crispr.mit.edu/). A pair of sgRNA oligos containing the guide sequence were annealed and ligated to the *Bbs*I-linearized vector pX330. The sgRNA oligos targeting *Hbxip*: 5’-CACCGTCTCACTATTCTCTAGGCCG-3’ and 5’-AAACCGGCCTAGAGAATAGTGAGAC-3’. The sgRNA oligos targeting *Tsc1*: 5’-CACCGGTGGTCAGGATGTGCAATAC-3’ and 5’-AAACGTATTGCACATCCTGACCACC-3’. The sgRNA oligos targeting *Lamtor3*: 5’-CACCGTCCTGTACAAAAAGTTGCCA-3’ and 5’-AAACTGGCAACTTTTTGTACAGGAC-3’. The sgRNA oligos targeting *Lamtor4*: 5’-CACCGTGTGCTGACCAACTCCGAGA-3’ and 5’-AAACTCTCGGAGTTGGTCAGCACAC-3’; 5’-CACCGCTGCTTGCTCATCGTTCTCA-3’ and 5’-AAACTGAGAACGATGAGCAAGCAGC-3’.

### Cell proliferation assay

1×10^5^ ESCs were plated in 12-well plates in triplicates. Cells were passaged continuously, and counted every 2 days.

### Western Blotting

ESCs were collected and dissociated in cold lysis buffer (20 mM HEPES, pH 7.7; 5 mM KCl; 1.5 mM MgCl2; 2 mM DTT; 2 mM PMSF) for 30 minutes on ice. Then the cell lysate was centrifuged at 12,000 rpm, 15 minutes at 4 °C. Protein concentration was measured using BCA Protein Assay Kit (Beyotime). The samples were resolved with SDS-PAGE gel and transferred onto PVDF membrane. Then, the PVDF membrane was incubated in blocking buffer for 1 hour at RT, followed by incubating with primary antibodies (*α*-Hbxip, CST, 14633S; *α*-Nanog, Bethyl Laboratories, A300-397A; *α*-Oct4, Santa Cruz, sc-5279; *α*-Tubulin, Sigma, ASB4800087; *α*-S6K1, Sigma, SAB4502686; *α*-p-S6K1, Sigma, SAB4301518; *α*-H3, Abcam, ab1791) at 4 °C overnight. After washing three times, the membrane was incubated with HRP-liked secondary antibodies at RT for 1 hour. Immunoreactivity was detected by ECL Plus (Beyotime). Digital images were taken with by the automatic chemiluminescence imaging analysis system (Tanon).

### Co-immunoprecipitation (Co-IP)

ESC extracts were prepared using lysis buffer (20 mM Tris-HCl, pH 8; 137 mM NaCl; 10% glycerol; 1% NP-40; 2 mM EDTA, supplemented with protease inhibitor and PMSF). FLAG antibody (M2) conjugated magnetic beads were washed in TBS buffer (50 mM Tris HCl, 150 mM NaCl, pH 7.4) for three times. ESC extracts were incubated with 15 μl M2 beads at 4 °C overnight. The immunoprecipitates were washed three times using TBS buffer, eluted in 2× SDS-PAGE loading buffer, and subjected to mass spectrometric analysis.

### RNA purification and quantitative reverse-transcription PCR

Total RNA was purified using TRIZOL (Ambion) according to the manufacturer’s instructions. cDNA was synthesized using Reverse Transcription Kit (Roche). PCR reactions were performed with FastStart Universal SYBR Green Master (Roche) in a Bio-Rad IQ5 system. PCR cycling conditions were 95°C for 2 min, 40 cycles of 95°C for 15 s, 58°C for 15 s, and 72°C for 30 s, and then a melting curve of the amplified DNA was acquired. Quantification of target genes was normalized with β-Actin. Primers were listed in Table S1.

### RNA sequencing

Total RNA was purified using RNeasy Mini Kit (Qiagen) according to the manufacturer’s instructions. Construction of RNA sequencing library, sequencing with BGISEQ-500, and bioinformatic analysis were performed by BGI Tech (Shenzhen, China). The RNA-seq data set (accession number GSE104242) has been deposited in the Gene Expression Omnibus database.

### Alkaline phosphatase (AP) assay

After washing once with PBS, mouse ESCs were stained with alkaline phosphatase substrate kit III (Vector) according to the manufacturer’s instruction.

### Colony forming assay

Mouse ESCs were suspended in ESC medium and cultured in a 6-well plate (800 cells per well). After 4-6 days culture, ESCs were stained for AP. The AP-positive colony were counted under microscope.

### Embryoid body (EB) forming assay

Mouse ESCs were suspended in LIF free ESC medium (40,000 cells/ml) and cultured in 25 μl hanging drops. EBs were collected on day 4.

### Statistical analysis

All images were processed with Photoshop 2021 (Adobe) and Image J (National Institutes of Health). All data were analyzed using Microsoft Excel and Prism software (GraphPad Software). Statistical analyses were performed with *χ*^2^, Student’s *t* or two-way ANOVA test, which were indicated in the figure legends. Statistically significant p-values were indicated in figures as *, p < 0.05; **, p < 0.01; ***, p < 0.001. And ns (not significant) marks p-value> 0.05.

## Acknowledgments

L.C. was supported by the National Key R&D Program of China (Grant No. 2018YFA0107002 and 2018YFC1313003), the National Natural Science Foundation of China (Grant No. 31871485), the Natural Science Foundation of Tianjin (Grant No. 18JCJQJC48400), the 111 Project Grant (B08011), and the Fundamental Research Funds for the Central Universities. Y.Q. was supported by the China Postdoctoral Science Foundation (Grant No. 2019M660980) and the National Natural Science Foundation of China (Grant No. 32000600).

## Author Contributions

Y.Q., P.N., Q.Z., X.D., X.W. Z.Y. and L.W. performed experiments, Y.Q. and P.N. analyzed the data and contributed to the paper writing, L.Y. and L.C. designed the experiments and wrote the paper.

