## Supplemental materials for "Hbxip (Lamtor5) is essential for embryogenesis and regulates embryonic stem cell differentiation through activating mTORC1"

### Supplementary Figure Legends

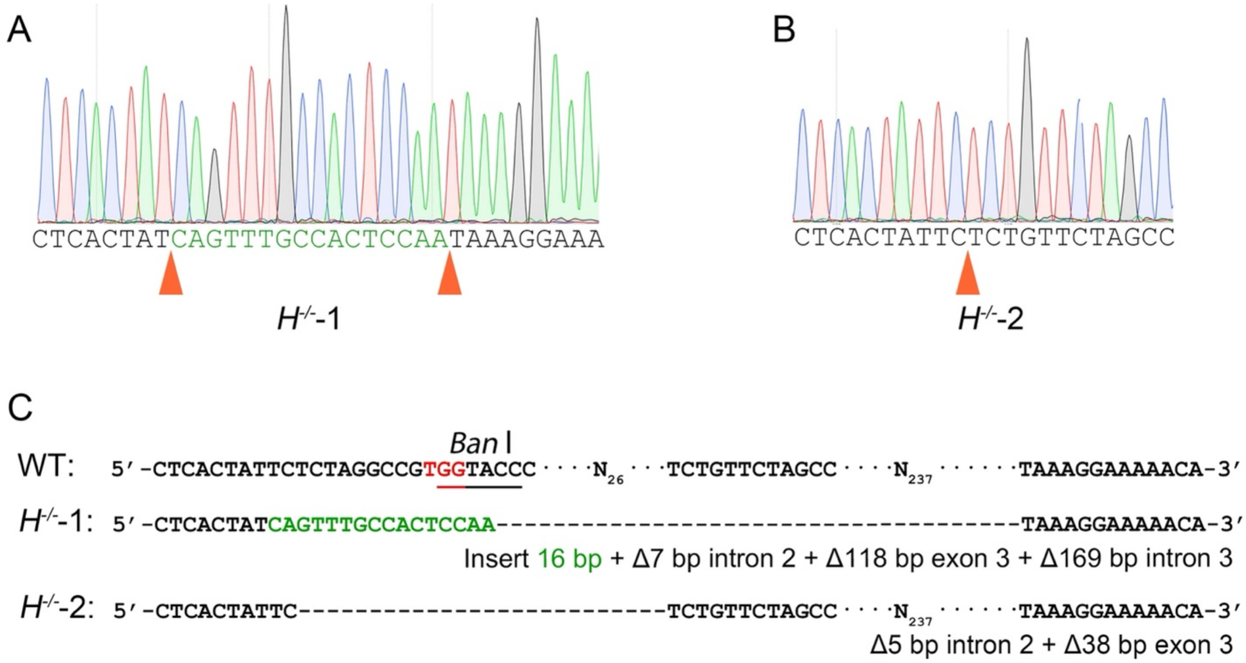


**Figure S1. Validation of *Hbxip*$^{-/-}$ ESC clones.** (Related to Figure 2)

(A) and (B) Sequencing chromatograph of the *Hbxip* gene around the Cas9 targeting site in $H^{-/-}$-1 (A) and $H^{-/-}$-2 (B) ESCs. Orange triangles mark the indel mutations introduced by Cas9 cutting.

(C) Sequence alignment of the *Hbxip* gene around the Cas9 targeting site in WT, $H^{-/-}$-1 and $H^{-/-}$-2 ESCs.

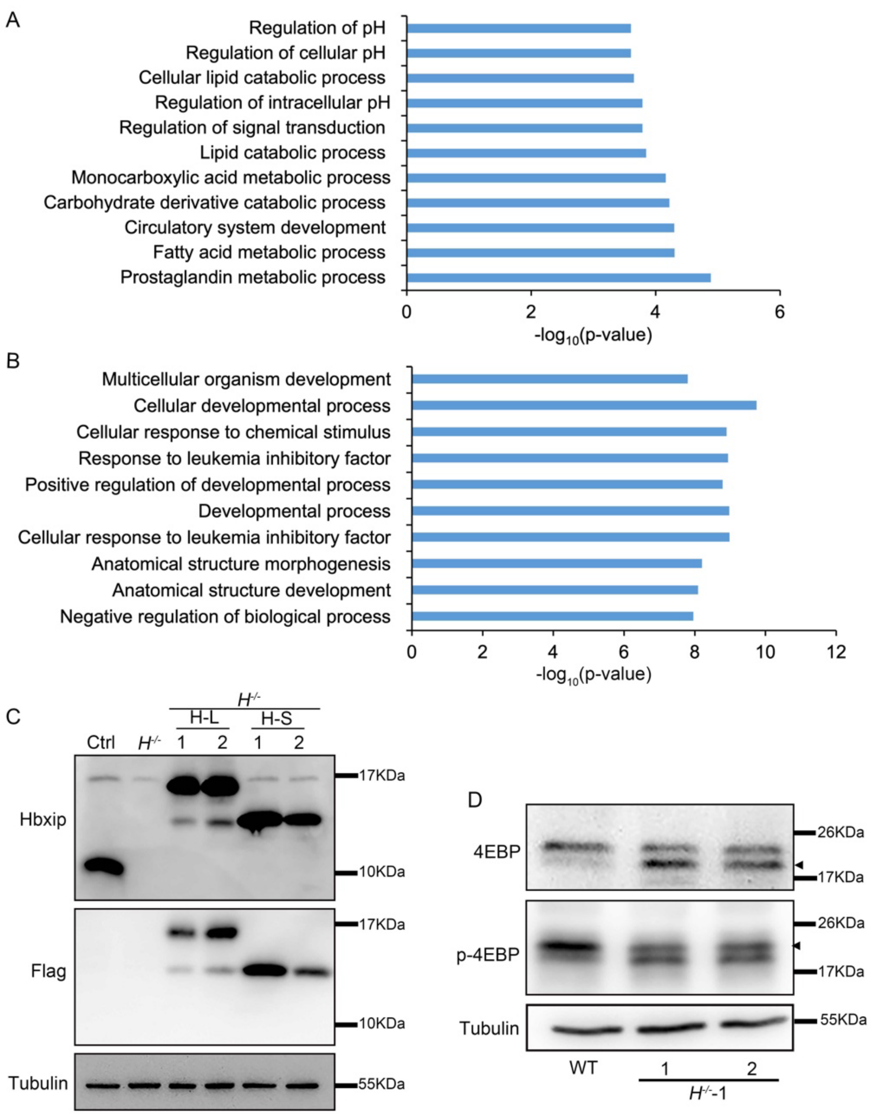


**Figure S2. *Hbxip KO* affects gene expression and mTORC1 activity.**

(A) GO annotation of the upregulated genes in *Hbxip*$^{-/-}$ ESCs (Related to Figure 2G and 2H). (B) GO annotation of the commonly upregulated genes by *Hbxip* KO (Related to Figure 3C-F). (C) Validation of the overexpression of Flag tagged Hbxip, including H-L and H-S (Related to Figure 3G). (D) Western blots of 4EBP and p-4EBP in WT and *Hbxip*$^{-/-}$ ESCs demonstrate the reduced mTORC1 activity in *Hbxip*$^{-/-}$ ESCs (Related to Figure 5D). Triangles mark the specific bands for 4EBP and p-4EBP. For Western blot, n=3.

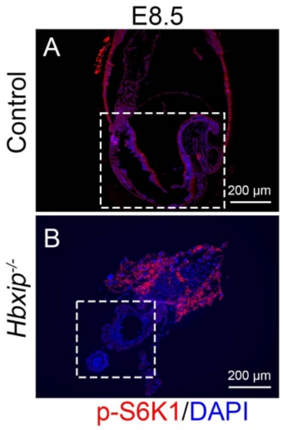


**Figure S3. Reduced mTOCRC1 activity in *Hbxip*$^{-/-}$ embryos.**

Immunofluorescence staining of p-S6K1 in control (A) and *Hbxip*$^{-/-}$ (B) E8.5 embryo sections. Dashed rectangles mark the embryo part. Control (including WT and *Hbxip*$^{+/-}$), n=4; *Hbxip*$^{-/-}$, n=4.

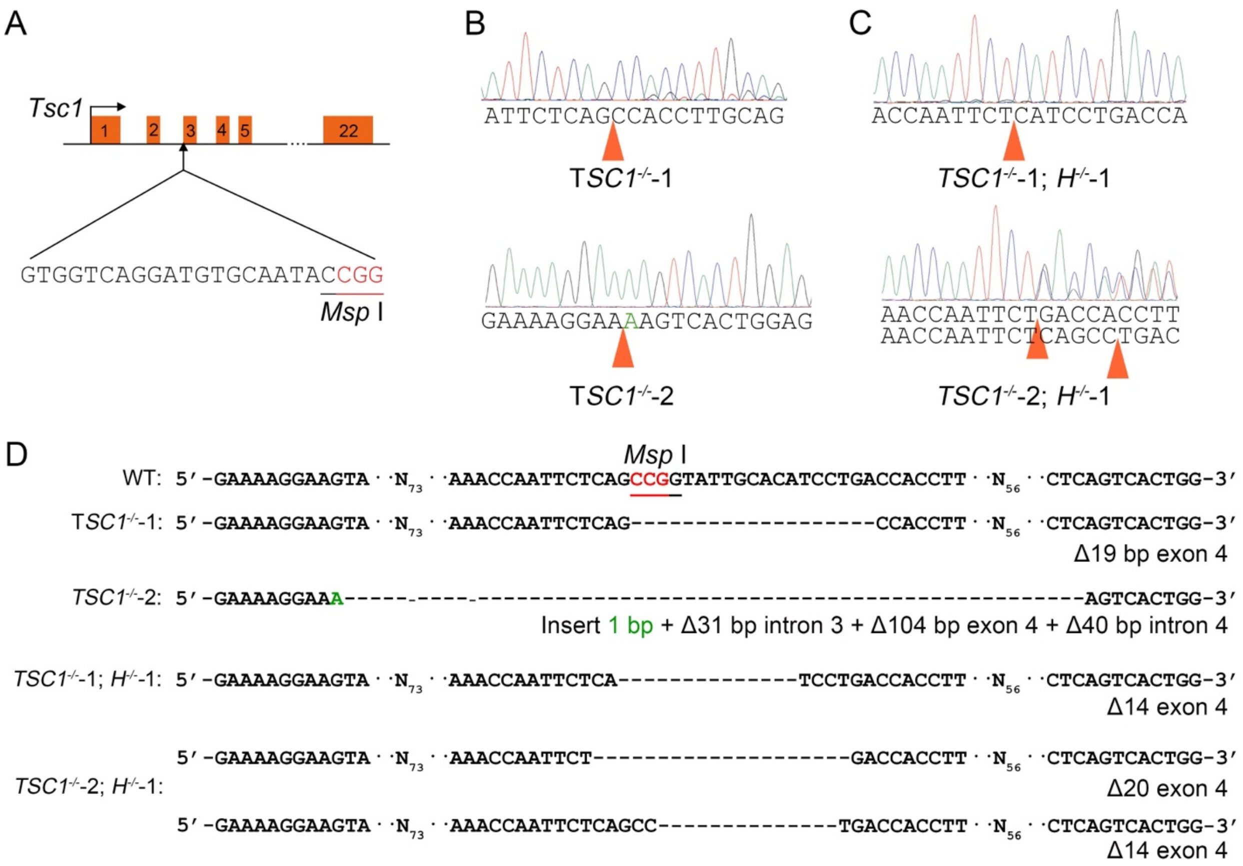


**Figure S4. Validation of *Tsc1*$^{-/-}$ ESC clones.** (Related to Figure 5)

(A) Schematic illustration of the strategy for knocking out *Tsc1* in ESCs. The targeting sequence of sgRNA is shown. The protospacer-adjacent motif (PAM) is highlighted in red, and the *Msp* I site is underlined. (B) and (C) Sequencing chromatograph of the *Tsc1* gene around the Cas9 targeting site in *Tsc1*$^{-/-}$ (B) and *Tsc1*$^{-/-}$; *H*$^{-/-}$-1 (C) ESCs. Orange triangles mark the indel mutations introduced by Cas9 cutting. (D) Sequence alignment of the *Tsc1* gene around the Cas9 targeting site in WT, *Tsc1*$^{-/-}$ and *Tsc1*$^{-/-}$; *H*$^{-/-}$-1 ESCs.

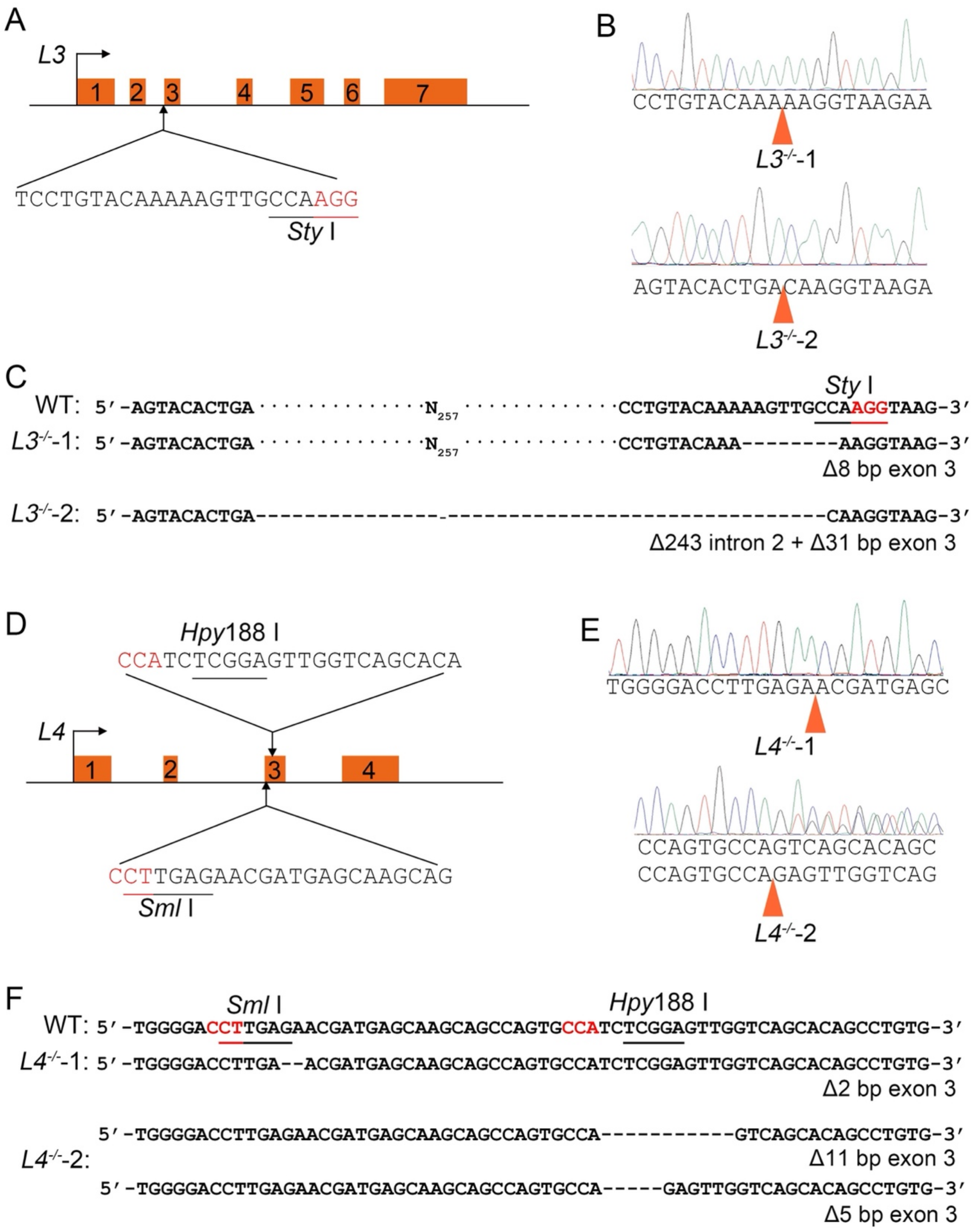


**Figure S5. Validation of *Lamtor3*$^{-/-}$ and *Lamtor4*$^{-/-}$ ESC clones.** (Related to Figure 6)

(A) and (D) Schematic illustration of the strategy for knocking out *Lamtor3* (A) and *Lamtor4* (D) in ESCs. The targeting sequences of sgRNAs (one for *Lamtor3* and two for *Lamtor4*) are shown.

The protospacer-adjacent motif (PAM) is highlighted in red, and the *Sty* I, *Sml* I and *Spy*188 I sites are underlined. (B) and (E) Sequencing chromatograph of the *Lamtor3* and *Lamtor4* gene around the Cas9 targeting site in *Lamtor3*$^{-/-}$ (B) and *Lamtor4*$^{-/-}$ (E) ESC clones. Orange triangles mark the indel mutations introduced by Cas9 cutting. (C) and (F) Sequence alignment of the *Lamtor3* (C) and *Lamtor4* (F) gene around the Cas9 targeting site in WT, *Lamtor3*$^{-/-}$ (C) and *Lamtor4*$^{-/-}$ (F) ESCs.

### Supplementary Tables

#### Table S1. Primers for quantitative RT-PCR

| Gene | Forward | Reverse |
|---|---|---|
| *Nanog* | TACAAGGGTCTGCTACTGAGATGC | TTGGGACTGGTAGAAGAATCAGGG |
| *Oct4* | ATCAGCTTGGGCTAGAGAAGGATG | AAAGGTGTCCCTGTAGCCTCATAC |
| *Sox2* | GCGGAGTGGAAACTTTTGTCC | CGGGAAGCGTGTACTTATCCTT |
| *Nestin* | CTGGATCTGGAAGTCAACAGAGGT | ATCCTCAGTTTCCACTCCTGTAGC |
| *Gata4* | GCTATGCATCTCCTGTCACTCAGA | CCAAGTCCGAGCAGGAATTTGAAG |
| *Gata6* | CTTCTCCTTCTACACAAGCGACCA | ATACTTGAGGTCACTGTTCTCGGG |
| *T* | CATCGGAACAGCTCTCCAACCTAT | TACCATTGCTCACAGACCAGAGAC |
| *Celsr2* | CACGATGGCCTGAGGGTTT | CCTTGTGGAGAAAGGTGTCCT |
| *Dlx3* | CACTGACCTGGGCTATTACAGC | GAGATTGAACTGGTGGTGGTAG |
| *β-Actin* | CAGAAGGAGATTACTGCTCTGGCT | TACTCCTGCTTGCTGATCCACATC |

#### Table S2. Differentially expressed genes (DEGs) identified in *Hbxip*$^{-/-}$ ESCs, compared to WT ESCs.

#### Table S3. DEGs identified in differentiated *Hbxip*$^{-/-}$ ESCs, compared to WT ESCs.

WT, *H*$^{-/-}$-1, and *H*$^{-/-}$-2 ESCs were differentiated for 4 days by two methods, LIF withdrawal and EB. RNA purified from these differentiated cells were subjected to RNA-seq analysis.

#### Table S4. Hbxip interacting proteins identified by co-IP and mass spectrometry.
